# Beyond the glass: testing visitor effects on the behavior of teleost fish in public aquaria

**DOI:** 10.64898/2026.09.17.752300

**Authors:** Lilou Vincent, Théophile Turco, Jerome Mourin, Joel Attia, Anne Sophie Tribot

## Abstract

Zoos and public aquariums serve as essential hubs for wildlife conservation, public education, and scientific research; however, the success of these missions depends upon maintaining high animal welfare standards despite constant human presence. A primary concern is the ‘visitor effect’: the modification of animal behavior due to human presence. While this phenomenon is well-documented in captive mammals and has been explored in Chondrichthyans, it remains largely unexamined in teleost fish, despite evidence of their sensitivity to human stimuli in natural environments. This study addresses this knowledge gap by investigating the behavioral responses of 13 teleost species to varying levels of visitor density, direct human solicitations, and ambient noise. Our multivariate analysis reveals that the total number of visitors present in the aquarium and the direct sonic human solicitations (such as knocking on the glass) are significant drivers of behavioral changes. We observed high inter-taxonomic variability: while *Euxiphipops sextriatus, Dascyllus aruanus*, *Zebrasoma flavescens*, *Chaetodontoplus mesoleucus* and *Pygoplites diacanthus* showed marked sensitivity through significant modifications in their behaviors the others (*Amphiprion percula, Chaetodon rafflesia, Chromis viridis, Chrysiptera parasema, Heniochus chrysostomus*, and *Lactoria cornuta*) showed no obvious behavioral changes. Furthermore, acoustic analysis indicates a complex soundscape where global visitor density correlates with increased sound amplitude, particularly in the 100 Hz to 400 Hz range, a critical band for teleost hearing, suggesting that visitor-induced noise is distinct from the aquarium’s technical background noise (pumps, aerators…). These findings underscore that the ‘visitor effect’ is not a uniform phenomenon but a complex interaction shaped by species-specific sensitivity and distinct anthropogenic actions. Our results highlight the necessity of incorporating species-specific behavioral data and acoustic monitoring into welfare management protocols to better mitigate the impact of human presence on captive fish populations.

## INTRODUCTION

As part of their core mission, zoos and public aquariums are tasked with actively participating in conservation, scientific research, and environmental education. These obligations are reinforced by international frameworks, such as the *World Zoo and Aquarium Conservation Strategy* (WAZA, 2005), and regional regulations, including the EU Directive 1999/22/EC (EU Directive, 2016). For accredited institutions, these goals are not merely peripheral; both the *Association of Zoos and Aquariums* (AZA) and at our local scale the French “*Union des Conservateurs d’Aquariums”* (UCA) mandate that member institutions prioritize active conservation and scientific advancement, positioning these functions as the central pillars of their existence (Hutchins et al., 2019; Larson, 2017; Ripple et al., 2021; Teletchea et al., 2023). Furthermore, professional bodies like the IUCN SSC state that zoos/aquariums must integrate species management, field research, and social science to counter biodiversity loss (Correia et al., 2024; SSC-IUCN, 2023). However, balancing rigorous conservation imperatives with their traditional public recreation role creates an inherently contradictory dual mission (Davey, 2007). Consequently, constant human presence in these educational spaces presents significant animal welfare challenges (Hashmi and Sullivan, 2020; Sherwen and Hemsworth, 2019). High visitor density, in particular, acts as a primary stressor (Stevens et al., 2013), often accompanied by unpredictable environmental fluctuations, such as increased ambient noise and sudden, erratic human movements (Boyle et al., 2020; Quadros et al., 2014). Furthermore, direct attempts at interaction by visitors can further disrupt an animal’s perceived security and increase its overall stress levels (Sherwen et al., 2014).

These complex interactions are encapsulated by the “visitor effect,” a term popularized by Hosey (2000) to describe the multifaceted influence exerted by human visitors on the behavior, physiology, and overall welfare of captive animals. Research indicates that this phenomenon is not necessary negative; its effects can range from potential enrichment to significant distress, encompassing neutral behavioral responses (see the review of Sherwen and Hemsworth, 2019). These outcomes are mediated by a complex interplay of species-specific sensitivity, individual temperament, and enclosure design (Davey, 2007; Hosey, 2000; Sherwen and Hemsworth, 2019). The visitor effect is also driven by various anthropogenic factors, including visitor presence, density, proximity, and noise levels (Davey, 2007; Hashmi and Sullivan, 2020). For captive animals, negative human behaviors, such as banging on glass or shouting, are often correlated with increased attendance and can push animals beyond their tolerance thresholds, forcing them to retreat to out-of-stimuli areas when they exist (Collins et al., 2023; Fernandez et al., 2009). For instance, increased visitor density has been linked to a reduction in positive social interactions in cotton-top tamarins (*Saguinus oedipus oedipus*) (Glatston et al., 1984) and shift in behaviors including increased pacing in captive felids (Mallapur and Chellam, 2002; Suárez et al., 2017). Furthermore, the visitor effect is a multimodal process involving visual, auditory and olfactive disturbance (Clark and King, 2008; Rose and Rice, 2025). Morgan and Tromborg (2007) highlighted that increased ambient noise, which often scales with visitor numbers, leads to significant behavioral shifts, as exemplified by orangutans (*Pongo pygmaeus*) at Chester Zoo, which exhibited heightened vigilance in response to amplified background noise (Birke, 2002). The visitor effect is traditionally assessed through behavioral observation, focusing on both abnormal stereotypical behaviors (e.g., pacing) and the disruption of natural behaviors like feeding and reproduction (Queiroz and Young, 2018; Wells, 2005). However, studies are increasingly incorporating physiological measures, such as glucocorticoid levels, to provide a more comprehensive view of animal welfare (Davis et al., 2005; Sherwen et al., 2015). Recognizing these impacts, modern management strategies now advocate for exhibit designs that provide “off-show” areas for retreat (Hashmi and Sullivan, 2020). They also promote the implementation of ethical assessment protocols, such as the Animal–Visitor Interaction Protocol (AVIP), to balance human engagement with the need for animal security (de Mori et al., 2019).

Past research on the visitor effect remains heavily biased toward non-human primates and felines, leaving aquatic species, particularly Osteichthyans, including teleosts, significantly underrepresented (Boyle et al., 2020). This gap in knowledge may stem from a perceived lack of empathy for fish. According to Würbel (2009), human empathy is often based on perceived morphological and behavioral similarities, such as facial structure and the facial muscles that enable recognizable expressions, between humans and other animals.. This, in turn, influences the motivation to ensure animal welfare. Despite being frequently perceived as less responsive to anthropogenic disturbance than terrestrial wildlife, fish (Osteichthyans, Chondrichthyans, and Cyclostomes) are highly sensitive to their environment and actively modulate their behavior in response to external changes (Braithwaite and Salvanes, 2005). Evidence indicates that human presence often functions as a significant environmental stressor, frequently interpreted by fish as a predatory threat. This interaction triggers an immediate physiological stress response characterized by the activation of the hypothalamic-pituitary-interrenal (HPI) axis and increased cortisol levels (Geffroy et al., 2018). Chronic exposure to human visitation has been linked to profound maladaptive outcomes, including the suppression of natural territorial and reproductive behaviors, heightened flight initiation distances (Bessa and Gonçalves-de-Freitas, 2014; Samia et al., 2019), and long-term physiological costs such as hematological decline and altered gene expression (Geffroy et al., 2018; Semeniuk et al., 2009). Consequently, the human presence represents a critical, yet often underestimated, factor influencing the behavior, physiology, and welfare of fish populations across both natural and captive environments (Boyle et al., 2020).

While recent studies have begun to examine the impact of visitor density and interaction on the behavior of Chondrichthyans, such as sharks and rays, in captive settings (Boyle et al., 2020; Galante et al., 2026; Truax et al., 2023), these findings cannot be readily extrapolated to other aquatic animals. Research on Chondrichthyans has increasingly utilized behavioral and spatial proxies, such as activity budgets, ledge use, and the avoidance of high-traffic areas, to monitor welfare (Galante et al., 2026; Hart et al., 2022; Wosnick et al., 2023). These findings are not readily applicable to Osteichthyans (bony fish), which constitute the majority of aquarium biomass and diversity due to fundamental physiological, metabolic, and anatomical differences. Emerging evidence from the field confirms that bony fish are highly sensitive to anthropogenic stimuli, experiencing significant behavioral and physiological shifts in the presence of visitors (Bessa et al., 2017; Samia et al., 2019). Studies indicate that factors ranging from visual cues to direct physical interaction can trigger stress responses, including elevated cortisol levels and the suppression of natural behaviors (Barcellos et al., 2007; Lawrence et al., 2021). While these responses are context-dependent (Leong et al., 2009; Silva et al., 2025), around 60% of fish species exhibit detectable changes in response to visitors, highlighting the need for species-specific research (Boyle et al., 2020). Moving beyond generalized assumptions is critical to developing evidence-based welfare management for diverse teleost populations.

Extending the visitor effect framework to these species in captive environments is the next critical step; it would not only bridge a significant taxonomic gap in welfare science but also clarify the mechanisms of adaptation in diverse aquatic environments. Insights gained from such an approach are essential for developing evidence-based management strategies, including refined exhibit design and visitor management protocols, which aim to safeguard the welfare of these animals. To provide these necessary insights, this study investigated the impact of both visual and auditory visitor disturbances on a multi-species group of teleost fish at the Aquarium de Lyon (France). We distinguished three levels for the visitor effect on the fish behavior: total frequentation in the building, visitor density and human behaviors in front of the exhibit. Concerning the acoustic events, we considered the noise variations in the whole structure, due to the evolution of the total frequentation within a day and between days. We also consider the number of people passing in front of the exhibit (trampling sounds) and the sounds due to specific human behaviors in front of the exhibit: jumping, touching the glass, *etc.* Using this multi-sensory approach, this study aims to determine the extent to which total frequentation in the building, visitor density and human behaviors in front of the exhibit influence the behavioral responses of teleosts.

Our primary hypothesis is that high visitor total frequentation/high visitor density facing the exhibit and active human behaviors act as visual and acoustic stressors, inducing behavioral shifts in fish such as an increased reliance on refugia or reduced activity levels. Conversely, we hypothesize that periods of low visitor density will correlate with an increase in exploratory behavior and the expression of natural agonistic interactions.

## MATERIAL AND METHODS

### Study subjects and captivity conditions

This study was conducted on a triangular (3 x 3 x 2.75 m) exhibit named “*La Nouvelle-Calédonie*” (figure 1) at the Aquarium de Lyon (France). The exhibit houses 23 wild-caught teleost fish representing 13 species (Table 1), alongside a colony of *Echinopora lamellosa* (1.6 m³) and other corals. All specimens were supplied by Nguyen International (Kingersheim, France). Data collection was carried out in May 2025, 30 months after the introduction of the last fish (Table 1); population interactions can be considered relatively stable.

**Figure 1:**
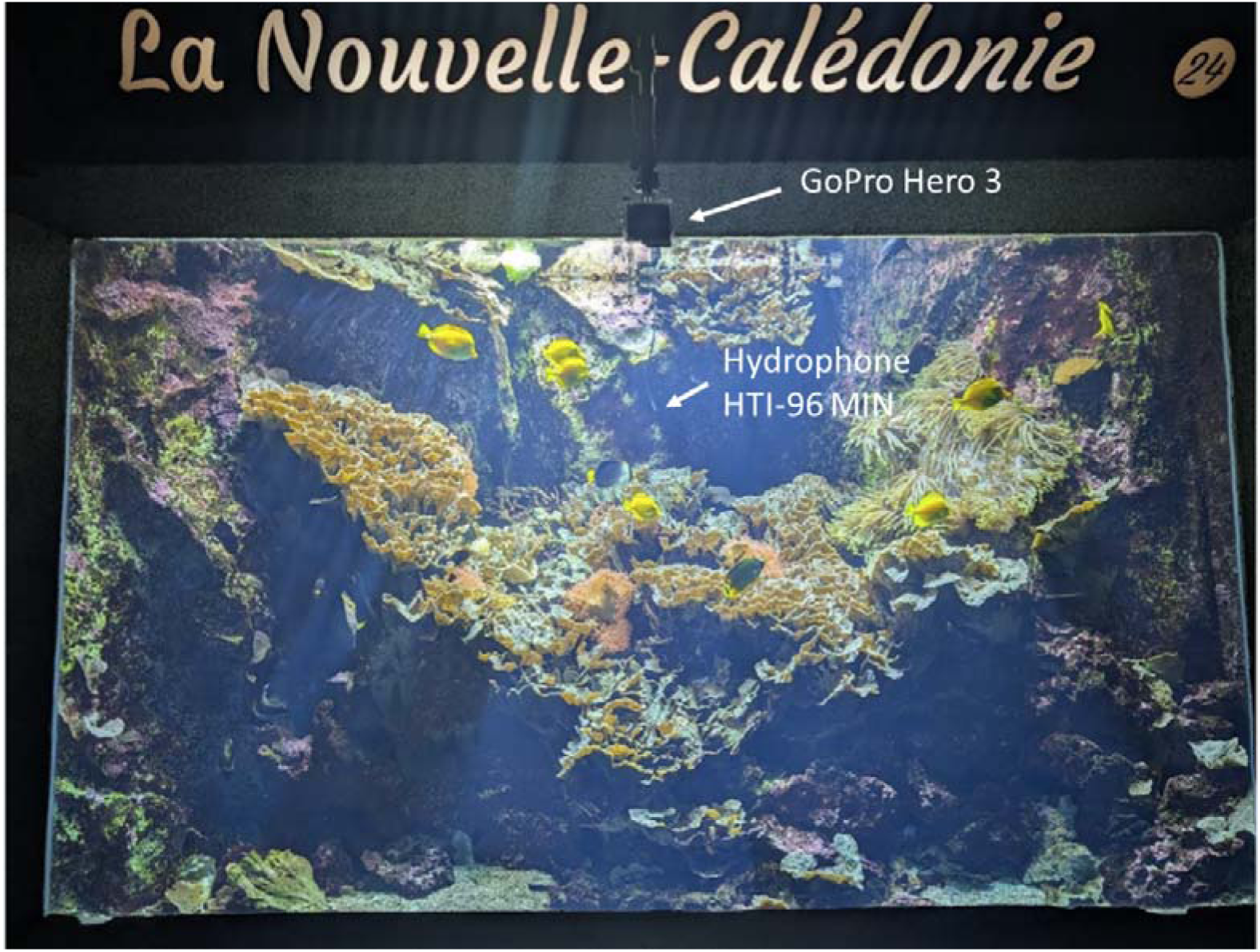
Video and acoustic recording system at Aquarium de Lyon. Overview of exhibition tank #24 (“New Caledonia”), home to a community of 23 fish. The positioning of the external GoPro Hero 3 camera (mounted high up) and the HTI-96-MIN hydrophone (submerged for acoustic recordings) is highlighted. CC Lilou Vincent.

**Table 1:** Taxonomy and behavioral characteristics of the studied teleost species.

| Species | Scientific family | Date of arrival | Nb of individuals | Territorial | Sociality |
| --- | --- | --- | --- | --- | --- |
| *Acanthurus achilles* | Acanthuridae | 09/2003 | 1 | Yes | Scattered group |
| *Amphiprion percula* | Pomacentridae | 08/2015 | 2 | Yes | Colony |
| *Chaetodon rafflesii* | Chaetodontidae | 06/2017 | 1 | No | Pair |
| *Chaetodontoplus mesoleucus* | Pomacanthidae | 10/2006 | 1 | No | Group |
| *Chelmon rostratus* | Chaetodontidae | 01/2017 | 1 | Yes | Pair or solitary |
| *Chromis viridis* | Pomacentridae | 10/2012 | 2 | Yes | School |
| *Chrysiptera parasema* | Pomacentridae | 01/2013 | 1 | Yes | Pair or solitary |
| *Dascyllus aruanus* | Pomacentridae | 10/2002 | 1 | Yes | School |
| *Euxhiphops sexstriatus* | Pomacanthidae | 10/2020 | 1 | Yes | Pair or solitary |
| *Heniochus chrysostomus* | Chaetodontidae | 12/2014 | 2 | No | Pair |
| *Lactoria cornuta* | Ostraciidae | 02/2023 | 1 | No | Solitary |
| *Pygoplites diacanthus* | Pomacanthidae | 09/2015 | 1 | No | Pair or solitary |
| *Zebrasoma flavescens* | Acanthuridae | 04/2017 | 8 | No | Group or solitary |

Observations were conducted *via* a single viewing window composed of double-laminated tempered safety glass, consisting of two 19 mm-thick panes. The exhibit functions on a closed-circuit system, employing a surface overflow to facilitate water drainage, thereby ensuring a complete volume renewal is achieved every 48 hours. A circulation pump facilitates the circulation of water with a capacity of 32 m³/h. In order to minimize stress, environmental conditions are strictly regulated. Physico-chemical parameters are maintained at constant levels, with water temperature ranging from 27 to 28°C, salinity remaining at approximately 35.6 g. L ¹ and pH held at 8.1. Dissolved oxygen concentrations are consistently higher than 100% saturation. In order to sustain a naturalistic diurnal cycle, the photoperiod is configured to 13 hours of light and 11 hours of darkness, with a programmed gradual transition occurring at 08:30 and 19:30. The maximal intensity of light within the tank is 1,586 lux, in contrast to the significantly lower ambient light level of 3.05 lux maintained in the designated visitor area.

The nutritional requirements of the group under study were met through a set feeding regime, whereby the subjects were fed four times weekly, on Monday, Wednesday, Thursday, and Saturday, between 09:00 and 10:00. The diet consisted of spinach, crushed mussels, chopped shrimp, and salad, with the quantities of each component being approximately 20 g. In addition, a daily supplement of *Artemia nauplii* and zooplankton was administered every evening, with the exception of scheduled feeding days.

The selection of this particular exhibit was made on the basis of the presence of highly charismatic species, such as *Amphiprion percula* and *Lactoria cornuta* (Langlois et al., 2022), which guarantee high visitor attraction and sustained attendance around the tank. Furthermore, this exhibit houses a community characterized by a wide diversity of biological, behavioral, and ecological traits, ranging from solitary to schooling species (see Table 1), and varying in territoriality and dietary regimes. This multi-species composition provides an ideal baseline to explore how visitor affect teleosts according to their specific ecological profiles.

### Behavioral studies

#### Procedure for tracking fish behaviors

Twenty-minute recordings were conducted twenty times over a one-month period at the same times: 10:30, 11:30, 12:30, 14:20, 15:20, and 16:20, to capture the fish behaviors. Each recording was captured using a GoPro Hero 3 camera mounted in an elevated position. The camera was positioned exactly 60 cm from the viewing window (Figure 1). To ensure full coverage of the environment, the camera utilized its native ultra-wide field of view *via* its integrated, factory-standard f/2.8 6-element aspherical glass lens. Video data were captured at a resolution of 1080p/1920×1080 at a frame rate of 30 fps, encoded using the H.264 codec, and saved in .mp4 format. This wide-angle configuration enabled the simultaneous capture of the entire exhibit and the immediate visitor area directly in front of the viewing window. To minimize novelty effects, the subjects were habituated to the camera’s presence for one week prior to the start of the study. Activating the camera required walking past the exhibit and could disturb the fish’s movement. Therefore, video exploitation only began ten minutes after the camera was activated, to allow the fish time to settle. Similarly, to prevent any interference from staff, access to the back of the exhibit was prohibited 5 minutes before the recording activation and remained so until its end.

#### Procedure for tracking human behaviors

To evaluate human-animal dynamics, we quantified visitor presence across three cascading scales of resolution: total frequentation in the overall facility, visitor density facing the experimental exhibit, and fine-grained human behavior facing the exhibit. Total frequentation was recorded at the conclusion of each session to estimate total longitudinal exposure, operating under the assumption that all visitors transited past the exhibit at least once. This macro-level metric was complemented by local scale counts of local visitor density, which we quantified by recording the number of individuals within a 3.4-meter radius of the tank with both feet were positioned inside the radius, irrespective of their body orientation. Moving from spatial proximity to direct interactions, we monitored the specific behaviors of visitors facing the aquarium.

Following Hosey (2005) assertion that presence alone functions primarily as a baseline condition rather than a predictive variable, our analysis encompasses visitor active stimuli. We specifically targeted direct physical interactions at the glass interface, such as placing hands on or tapping the glass, as well as jumping or climbing over the low barrier in front of the aquarium. This tracking is critical because, although the radical refractive index mismatch at the flat glass wall renders the teleost eye severely hyperopic and causes the external world to appear structurally blurred (Douglas and Crawford, 2001), fish remain highly sensitive to outside actions. Even as light refraction compresses the aerial environment into a tight 97° Snell’s window and turns the rest of the glass barrier into a mirror (Dunn and Fitzgerald, 2020; Land and Nilsson, 2012), the teleost visual system is highly optimized to process whole-field visual motion rather than fine static details (Wang et al., 2020). Consequently, erratic visitor movements create distinct moving contrasts that downstream tectal circuits readily detect. Because aquatic animals can recognize individual faces, differentiate human roles, and significantly alter their swimming patterns or spatial positioning when humans approach the glass (Miller et al., 2023), visitor induced visual stimuli could act as powerful environmental disruptors that can directly skew behavioral data if left unmonitored. Because behavioral responses to anthropogenic exposure are highly species- and individual-specific (Boyle et al., 2020), visitor-fish interactions were documented on an individual basis whenever feasible. To preserve the integrity of these public behaviors and minimize observer bias, all sessions were conducted by a single, unobtrusive observer in plain clothes who maintained public interactions at a strict minimum and refrained from enforcing facility regulations regarding noise or flash photography.

### Fish and human behavioral analysis

#### Behavioral tracking

Fish and human behavioral data were analyzed using the software BORIS (version 8.27.10) (Friard and Gamba, 2016). The behavioral repertoire of the fish was categorized as follows:

- *Aggression*: Defined as the sum of chasing, biting, intimidation, and caudal extension.
- *Stress coping*: Defined as the sum of color changes, avoidance, fleeing, and refuge usage.
- *Exploration*: Defined as the sum of passing through, surface activity, and total spatial switches between zones covering depth levels (3 levels) and orientation (3 levels).
- *Feeding*: individual measurements for feeding.
- *Inactivity*: individual measurements for inactivity

The human behaviors were categorized in a single category encompassing tapping on the glass, adhering hands on the glass, and climbing the small wall in front of the exhibit.

The analysis involved counting of these behaviors throughout each 20-minute recording session. This methodology was applied to all individuals within the tank, with the exception of the seven *Zebrasoma flavescens* that could not be individually distinguished. For this species, observations were conducted at the group level; during each session, one individual was randomly selected to represent the cohort. In instances where two individuals from same species were present simultaneously into the exhibit, their data were merged into a single pooled. In total, the dataset comprised approximately 197 hours of video recordings, enabling a comprehensive assessment of the subjects’ behavioral responses across varying levels of visitor presence and acoustic stimuli.

#### Statistical processing

The data statistical analysis was performed using the software R (R Core Team, 2022) via RStudio. First, to visually explore the behavioral difference of each fish according to total frequentation in the building and human behaviors, we performed a Uniform Manifold Approximation and Projection (UMAP) using umap package (v0.2.10.0, Konopka, 2018). UMAP is a non-linear dimensionality reduction technique that excels at preserving both local and global data structures, allowing us to map high-dimensional behavioral profiles—comprising aggression, stress coping, exploration, feeding, and inactivity—onto a two-dimensional plane (Figure 2a). Prior to the UMAP construction, behavioral counts were centered and scaled (mean = 0, SD = 1) to account for differences in measurement units. We optimized the UMAP projection using the Euclidean distance metric, setting n_neighbors to 5 to restrict the neighborhood size—thereby capturing highly localized data relationships—and min_dist to 0.5 to allow a more dispersed, continuous distribution of points in the embedded space. This visualization enabled the identification of distinct behavioral clusters and facilitated the assessment of fish behavioral shifts relative to total public frequentation, visitor density in front of the exhibit and human behaviors. To statistically assess the multivariate behavioral responses of the teleost community, we fitted Multivariate Generalized Linear Models (ManyGLM) within the mvabund R package (v4.2.1; Wang et al., 2012), utilizing a negative binomial error structure to account for overdispersion. Unlike distance-based approaches such as PERMANOVA—which collapse multivariate data into a dissimilarity matrix and can confound true centroid shifts across time with within-group dispersion effects—this model-based framework fits GLMs directly to raw count data. By explicitly modeling the mean-variance relationship, this approach effectively handles the heteroscedasticity typical of behavioral datasets while providing robust, variable-specific statistical inference over time.

**Figure 2:**
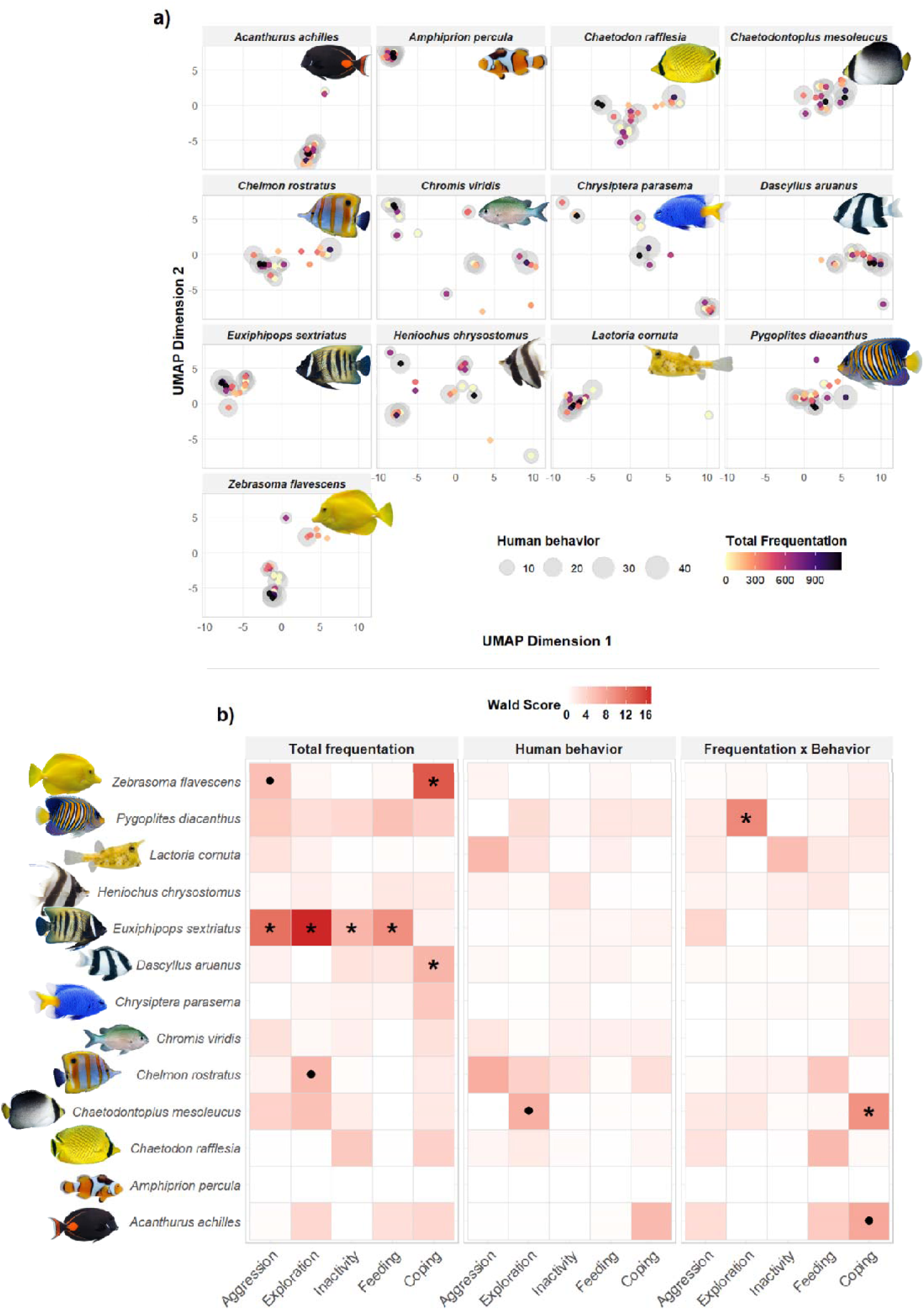
Fish behavioral response to total frequentation and human behavior. a) UMAP projection of behavioral states across aquarium species in relation to total frequentation. Each panel represents a distinct species. The behavioral space was constructed using UMAP dimensionality reduction on composite behavioral scores (aggression, exploration, feeding, inactivity and stress coping). Each point represents a single individual during a unique observation period, color-coded represent the total frequentation in the structure while the size of grey shaded circles indicates the presence and magnitude of visitor behavior. The proximity of points on the UMAP axes reflects the similarity of behavioral profiles within each species. b) This heatmap illustrates the behavioral responses of different fish species to total frequentation, human behavior, and their interaction (Frequentation x Behavior”). The color intensity represents the Wald Score, indicating the strength of the impact. Statistical significance and trends are marked as follows: Star (*): Indicates a statistically significant effect (p < 0.05) or Point (•): Indicates a behavioral trend (0.05 < p < 0.10).

Model selection was performed by minimizing the total sum of the Akaike Information Criterion (AIC) across all candidate models, evaluating potential interactions among visitor density (passersby facing the exhibit), total frequentation (overall building occupancy), human behavior, and fish species. While we also evaluated the effect of visitor density in front of the exhibit, models incorporating this parameter underperformed. Ultimately, the top-performing model retained the interaction between total frequentation and human behavior, alongside the additive effect of fish species (Total Frequentation × Human Behavior + Species). Statistical significance was assessed using 999 iterations of PIT-trap resampling. To identify specific behavioral drivers, we calculated adjusted univariate p-values for each ethogram category using a step-down multiple testing procedure.

To identify species-specific behavioral responses, the global model was partitioned by Species. We calculated univariate Wald statistics and adjusted p-values for each behavioral category within the ethogram (aggression, exploration, inactivity, feeding, and stress coping). Multiple testing was accounted using a step-down resampling procedure to control the family-wise error rate. The resulting behavioral shifts were visualized using a comparative heatmap framework, faceted by experimental effect (Total Frequentation, Human Behavior and their interaction).

### Soundscape study

#### Recording set up and procedure

To characterize the aquarium’s soundscape variation due to the total frequentation, the visitor density in front of the aquarium and the human behaviors, we established an ambient baseline by conducting recordings prior to the aquarium’s daily opening, in the absence of visitors.

Then three recordings were collected during public operating hours to identify sound level fluctuations and spectral variations associated with public attendance, using a dual-sensor array to capture simultaneous underwater and airborne sound. During these three periods, visitor densities in front of the aquarium reached 23, 19, and 7 individuals, corresponding to total building frequentations of 294, 418, and 654 individuals, respectively.

Finally, to characterize the acoustic signatures of human behaviors, we performed simulation tests in the absence of visitors. The same operator tapped the glass with a closed fist or open palm at a consistent force, applaused, jumped, and ran, at varying distances from the aquarium.

Underwater sound pressure was recorded using a High Tech, Inc. (HTI) 96-MIN hydrophone (sensitivity −165 dB re. 1V µPa−1, flat frequency response from 2 Hz to 30 kHz), while airborne acoustic levels were monitored using a dedicated SMX-II microphone (sensitivity - 35 dBV/Pa, frequency response 20 Hz to 20 kHz). Both sensors were connected to a Song Meter (SM2+, Wildlife Acoustics Inc., Concord, MA, USA) acoustic recorder. All data were acquired at a sampling rate of 96 kHz and a 16-bit depth. Prior to the study, the entire recording chain was calibrated using a hydrophone (Bruel & Kjaer 8104, Naerum, Denmark; sensitivity −205 dB re. 1V μPa−1; frequency response from 0.1Hz to 180kHz) connected to a sound level meter (Bruël & Kjaer 2238 Mediator, Naerum, Denmark) to ensure accurate sound pressure level (SPL) measurements. The hydrophone was positioned at the rear central area of the tank to capture representative ambient sound while minimizing visual interference with the subjects.

#### Acoustic data processing and analysis

Acoustic analyses were performed in the R statistical environment using custom processing scripts using seewave (v2.2.3, Sueur et al., 2008) and tuneR (v1.4.7, Ligges et al., 2013) R packages. Because raw digital audio files record relative amplitude rather than absolute physical sound pressure, a calibration protocol was implemented to convert the digital signal into physical units. A fixed calibration offset was determined by anchoring the total energy of a baseline recording to a known reference level of 139.65 dB re. 1 µPa. For each acoustic recording, spectral characteristics and root-mean-square amplitudes were computed via a Short-Time Fourier Transform (STFT) utilizing a Hann window (window length = 4096 samples, 0% overlap). The derived calibration offset was applied uniformly across the computed frequency spectrum to yield absolute sound pressure levels. Broad-band acoustic levels are reported as the equivalent continuous sound pressure level (L_eq_) expressed in dB re. 1 µPa.

To examine the temporal stability of the soundscape, the three 8-min recordings were segmented into 10-second intervals. For each interval, the L_eq_ was extracted to create a time-series dataset. The resulting datasets were aggregated to calculate mean spectra and the corresponding minimum-to-maximum amplitude ranges (spectral envelopes) in order to make comparison of intra-aquarium soundscape variation according to total frequentation and visitor density in front of the aquarium. These profiles were plotted against the generally accepted hearing range for teleost fish. We also assessed the synchrony between the public noise and the intra-aquarium soundscape by calculating Pearson correlation coefficients. To ensure the reliability of the intra-aquarium noise analysis, time-series outliers exceeding 1.5 times the IQR were systematically eliminated. These outliers corresponded to transient events such as fish touching the hydrophone.

To evaluate the acoustic perceptibility of simulated human behaviors, we calculated the Signal-to-Noise Ratio (SNR). The SNR was computed by comparing the sound pressure levels of specific human-generated noise against the ambient background noise recorded around the events. We defined the temporal duration of each event as the interval during which the acoustic energy of an event remained at least 1.5 times higher to that of the noise, centered on the signal’s peak amplitude.

### Ethical norms

This study involved exclusively non-invasive behavioral observations of teleost fish under routine aquarium conditions, requiring no manipulation or disturbance of the animals. The facility in which the observations were conducted complies with the provisions of the French decree of March 25, 2004, regarding the protection of animals in captivity establishments (French Environmental Code). Regarding visitor’s data, human behavior was observed within a publicly accessible aquarium in strict accordance with European and French data protection regulations (GDPR and the French Data Protection Act). Because the observations were entirely non-intrusive and anonymous, no informed consent was required.

## RESULTS

### Effects of total frequentation in the building and human behaviors on fish behaviors

The multivariate analysis revealed that the overall behavioral repertoire of the fishes was significantly influenced by human total frequentation in the building (Wald = 3.35, p = 0.006) and the interaction between total frequentation and human behaviors (Wald = 5.21, p = 0.002). Furthermore, significant inter specific differences were observed, with species identity acting as a primary driver of behavioral variance (Wald = 32.66, p = 0.001). Univariate tests further confirmed that the fish species consistently accounted for significant differences across all behavioral categories (feeding, inactivity, aggression, stress coping, and exploration; all p < 0.001). Human total frequentation showed a significant negative relationship on stress coping (β = −0.175, Wald = 2.486, p = 0.017) and exploration (β = −0.562, Wald = 0.439, p = 0.017), indicating that higher public frequentation decreases these behaviors. Conversely, feeding, inactivity, and aggression did not show a statistically significant univariate response to total public alone. Finally, the interaction between human total frequentation and human behavior levels did not reach statistical significance at the univariate level for any of the individual behavioral categories (all p > 0.05).

When examined at the species level (Figure 2), sensitivity to human total frequentation and visitor behaviors exhibited high inter-taxonomic variability. *Euxiphipops sexstriatus* showed the most pronounced response, marked by significant decreases across its entire behavioral repertoire in response to total frequentation, including reductions in feeding (β = −1.785, Wald = 9.57, p = 0.020), inactivity (β = −5.792, Wald = 6.33, p = 0.030), aggression (β = −9.307, Wald = 12.09, p = 0.010), and exploration (β = −1.612, Wald = 17.16, p = 0.010). Similarly, *Dascyllus aruanus* and *Zebrasoma flavescens* experienced significant declines in stress-coping behaviors under high total frequentation (β = −7.355, Wald = 6.45, p = 0.030 and β = − 1.022, Wald = 13.95, p = 0.010, respectively), while *Z. flavescens* additionally exhibited a trend toward increased aggression (β = 1.904, p = 0.069). The interaction between human total frequentation and visitor behaviors significantly affected *Chaetodontoplus mesoleucus*, which displayed a reduction in its stress-coping response (β = −3.126$, Wald = 9.35, p = 0.020), and *Pygoplites diacanthus*, which showed an enhancement in exploratory behavior (β = 2.507, Wald = 10.14, p = 0.030). Furthermore, *C. mesoleucus* exhibited a marginal trend toward increased exploration under the interactive effect of frequentation and visitor behavior (β = 6.512, p = 0.069). Finally, weaker behavioral trends included a marginal reduction in stress coping for *Acanthurus achilles* (β = −4.345, p = 0.089) and a marginal decline in exploratory activity for *Chelmon rostratus* in response to total frequentation (β = −8.463, p = 0.08). Conversely, *Amphiprion percula*, *Chaetodon rafflesia*, *Chromis viridis*, *Chrysiptera parasema*, *Heniochus chrysostomus*, and *Lactoria cornuta* maintained high behavioral stability, displaying no significant shifts or trends across all tested variables (p > 0.10).

### Acoustic results

The analysis of the underwater acoustic environment reveals no correlation between noise level in the exhibit room and noise recorded within the exhibit across the three sampled time periods (Figure 3a). Instead, the correlation analysis yielded mixed results: one session showed a robust, significant negative correlation (r = −0.672, p < 0.001), while the two other sessions produced negligible coefficients (r = −0.169 and r = 0.010, respectively). These results demonstrate that the exhibit’s acoustic environment is not a simple linear function of aggregate ambient visitor noise, revealing a more nuanced interplay between the soundscape and human activity. Conversely, a clear relationship exists between total frequentation in the building and underwater noise levels (Figure 3b). Assuming the noise from technical systems of the aquarium is constant across all recording sessions, the observed variations in acoustic levels are attributable to the noise produced by the total number of visitors, likely resulting from vibrations transmitted to the aquarium through the facility infrastructure. As demonstrated by the spectral analysis, increased public attendance corresponds to higher mean sound levels across a broad frequency spectrum. This elevation is most pronounced within the 100 Hz to 400 Hz range, a critical band that encompasses the peak hearing sensitivity for many teleost species. However, some specific human behaviors, tapping with a closed fist and an open hand, and jumping, produced distinct acoustic signatures that emerged significantly above the background noise floor, with mean signal-to-noise ratios (SNR) of 22.4 dB, 25.7 dB, and 8.8 dB, respectively (Sup.mat. 1). These data validate a direct mechanism for quantifying human-animal acoustic interactions, illustrating that human behaviors can produce significant, albeit occasional, deviations from the exhibit’s baseline soundscape.

**Figure 3:**
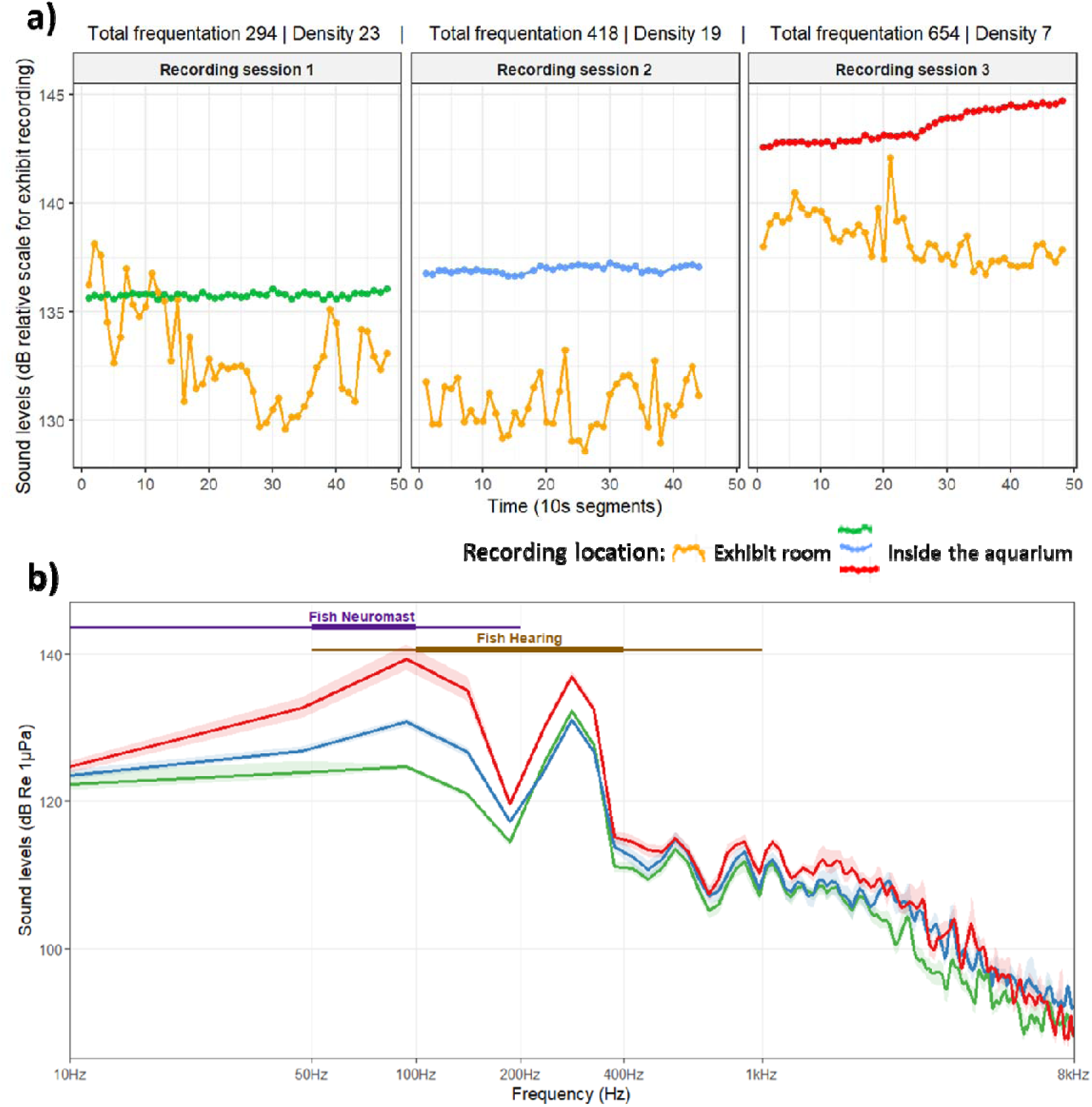
Acoustic analysis profiles. a) Acoustic synchrony analysis by session: time-series plot comparing ambient sound levels (dB) between the exhibit room (orange line) and the tank across three distinct recording sessions. Each data point represents the average amplitude calculated over 10-second intervals. “Total frequentation” and “Density” metrics are provided for each session to correlate visitor traffic with acoustic variance. The green, blue, and red lines correspond to exhibit recordings for Session 1, Session 2, and Session 3, respectively. b) Spectral analysis of sound levels: Frequency-dependent sound pressure levels (dB re 1µPa) across the observed spectrum (10 Hz – 8 kHz). Colored lines correspond to the respective sessions from panel *(a). Horizontal bars at the top of the plot indicate the physiological frequency ranges for specific fish sensory systems: the purple bar represents the range sensitive to the* f*ish* n*euromast*s *(lateral line system), and the brown bar represents the range relevant to* f*ish* h*earing*.

## DISCUSSION

These results support the hypothesis that the ‘visitor effect’ induces species-specific behavioral alterations, thereby modifying the community-level activity budgets of teleosts in public aquariums. Specifically, our data indicate that increased visitor total frequentation and associated human behavior drive a significant reallocation of time within these behavioral budgets. In this public aquarium, the “visitor effect” induces clear, species-specific behavioral shifts rather than a uniform community response. Species identity acted as the primary driver of behavioral variance, showcasing high inter-taxonomic variability ranging from highly sensitive species (e.g., *Euxiphipops sexstriatus* displaying alterations across its entire repertoire) to behaviorally stable species (e.g., *Amphiprion percula* and *Chromis viridis*). Historically, research on human presence effects in zoos and aquariums has struggled to reach a consensus, as observations often fluctuate between the perception of visitors as sources of chronic stress and their potential function as novel environmental stimulants.

Considerable evidence supports the stress-mediated hypothesis. Studies have demonstrated that exposure to typical aquarium stressors, such as acoustic disturbances (e.g. knocking on glass) and intense visual stimuli (e.g. photo-flashes), can induce physiological responses in fish, including elevated cortisol levels and suppressed metabolic rhythms (Leong et al., 2009). Furthermore, exposure to high-frequency tourism in natural or captive settings has been linked to long-term phenotypic shifts in stress-coping abilities, often resulting in increased avoidance behaviors and refuge usage (Geffroy et al., 2018). These “fight or flight” responses suggest that for many species, the exhibit may not be perceived as a safe enclosure, but rather a space permeated by threats (Lawrence et al., 2021). Conversely, an emerging body of literature challenges the notion that visitor presence is universally deleterious. Recent studies suggest that, under specific conditions, visitors may provide environmental enrichment. For example, research on various ray species and freshwater fish indicates that visitor presence can promote exploratory behaviors, increase foraging activity, and reduce periods of inactivity (Silva et al., 2025; Truax et al., 2023). Similarly, investigations into Atlantic cod have revealed “novelty-seeking” behaviors, where fish exhibit increased engagement with exhibit windows, seemingly attracted by the visual stimulus of the audience (Patton et al., 2023). Furthermore, the work of Fife-Cook and Franks (2021) with koi carp (*Cyprinius carpio*) underscores that some fish species are capable of forming positive human-animal bonds. In this paper, the fish seek for physical contact even after being fed to satiation, suggesting voluntary interest in human-animal interaction.

While previous research often quantifies this impact solely through measures of visitors facing the exhibit (Silva et al., 2025; Truax et al., 2023), our results highlight that this parameter is insufficient to assess the environmental changes perceived by fish. Instead, we show that visitor-induced disturbances in aquariums are complex, involving both visual and acoustic signals, at different scales: vibrations of the total human visitors walking in the whole structure, transmitted by the infrastructure facilities to the aquarium, sounds produced by specific human behaviors in front of the aquarium. Teleost species rely on hierarchical multisensory integration—simultaneously processing visual, acoustic, olfactory, and mechanical inputs—to inform critical life-history decisions such as foraging, predator avoidance, and social recognition (Derby and Sorensen, 2008; Moller, 2002; Ward and Mehner, 2010). In natural ecosystems, fish demonstrate remarkable behavioral plasticity, effectively navigating multisensory landscapes by switching modalities or prioritizing specific cues depending on the environmental context (Gardiner et al., 2014; Huijbers et al., 2012). In contrast, the exhibit environment imposes permanent environmental disturbances that can disrupt multisensory integration, leading to altered cognition and behavior in aquatic organisms (Kelley et al., 2018). The combination of visual movement—such as spectator locomotion—and percussive vibrations (e.g., tapping, jumping) creates a multimodal stressor that mirrors conditions known to force shifts in signaling modalities and risk assessment (de Jong et al., 2020; Landeira-Dabarca et al., 2019). Research demonstrates that exposure to human movement causes teleosts to over express refuge-seeking relative to foraging and territorial defense behaviors (Benevides et al., 2019; Pereira et al., 2016).

Because teleost species rely on sensitive auditory and lateral-line systems to interpret their environment, noise and mechanical vibrations transmitted through tank structures are readily perceived (Bart et al., 2001; Gutscher et al., 2011). Crucially, because different teleost lineages possess distinct sensory configurations, anatomical features, and ecological baselines, their susceptibility and response to these anthropogenic stimuli are highly variable stemming from a combination of biological factors and past experiences (Huntingford et al., 2006). This differential sensitivity need to be linked to the processes of habituation—a learned, stable decrease in response to repeated, neutral stimuli—or sensitization, defined as the amplification of a response (Coppola et al., 2013; Thompson and Spencer, 1966). While habituation is a biologically feasible mechanism for teleosts to manage neutral stressors (Radford et al., 2016; W. Wang et al., 2025), these underlying cognitive mechanisms remain poorly understood in captive environments (Köcher and Straumann, 2025). High visitor frequentation could preclude habituation by generating unpredictable, persistent stimuli without a viable escape route. These conditions could ultimately overwhelm sensory systems and hinder threat evaluation, thereby mirroring patterns of chronic disturbance seen in natural settings (Nanninga et al., 2017; Titus et al., 2015). Under these conditions, behavioral responses align with wild populations, where human presence is interpreted as a predatory threat, triggering increased flight initiation distances and heightened vigilance (Samia et al., 2019). Furthermore, the composition of the fish community represents a critical factor; because social context modulates adaptation styles (Castanheira et al., 2016; Frost et al., 2007), it is conceivable that the adaptive behaviors of less resilient species are disrupted not only by alterations in their direct social interactions, but also by behavioral “contamination”—the social transmission of stress or flight responses—stemming from the behavior of co-habiting species (Krause, 1993). Among the species observed, six did not change their behavior according to visitor presence: *Lactoria cornuta, Heniochus chrysostomus, Chrysiptera parasema, Chromis viridis, Chaetodon rafflesia*, and *Amphiprion percula*. The presence in this group of three Pomacentrids seems logical, as this family is widely documented for its exceptional hardiness and adaptability to environment (Nagelkerken et al., 2023; Olivotto and Geffroy, 2017). Their physiological resilience, including the ability to withstand wide fluctuations in water quality, might lead to the assumption of a taxonomic predisposition toward higher stress tolerance (Dhaneesh et al., 2012). However, this apparent absence of reaction to visitors could be attributed to successful habituation and the strategic use of secure micro-habitats, rather than an inherent lack of stress (Fakan et al., 2023).

Furthermore, relying solely on point-in-time visitor counts may fail to capture the cumulative intensity of exposure. As the duration and context of exposure are critical for welfare assessment (Lawrence et al., 2021), the accurate measurement of human impact requires detailed and long-term studies. Additionally, relying exclusively on fish behavioral metrics risks underestimating the impact on welfare. As documented in broader welfare studies, internal physiological stress often persists in the absence of observable behavioral changes (Millot et al., 2014; Rose, 2024). This disconnection between behavioral and physiological states occurs when animal’s coping mechanisms are masked or when the animals has reached a state of chronic, non-reactive stress (Rose, 2024). Consequently, observational methods alone may be insufficient for a complete welfare assessment. Future analyses must incorporate physiological markers, particularly cortisol commonly used as a stress indicator in teleost fish (Sadoul and Geffroy, 2019) to confirm whether these individuals are truly “tolerant” or merely exhibiting a suppressed behavioral response to human presence (Millot et al., 2014; Rose, 2024). Complementing this, Lawrence et al., (2021) identified that the physical architecture of an exhibit, such as the strategic placement of viewing barriers, acts as a primary mediator, allowing fish to modulate their exposure and stabilize their behavioral repertoire. These individuals consistently occupied zones furthest from the tank front, a spatial preference that likely indicates an underlying ‘shy’ personality type—a behavioral syndrome characterized by lower activity levels and a tendency to avoid risk (Blake and Gabor, 2014; Millot et al., 2014). Boyle et al., (2020) also demonstrated that individual variance is a critical factor, with responses differing significantly even within the same exhibit. This disparity in results—ranging from avoidance to attraction—highlights that the visitor effect is contingent upon a complex interplay of species-specific ecology, individual temperament, and exhibit design.

Our findings underscore the critical need for structural mitigation techniques, such as utilizing mechanical shock absorbers and selecting acoustic-dampening concrete or acrylic walls (Davidson et al., 2007) when designing aquarium (Anderson, 2013; Lu et al., 2025; Rose and Rice, 2025). Specifically, our study highlights that sound could propagates into aquarium through structural pathways, and seems heavily influenced by the facility’s architecture. Literature also reports that noise from air born sources of adjacent public areas—such as banquet halls, tunnels, and guest viewing rooms—transmits directly through aquarium boundaries, including acrylic and glass viewing windows (Jackson et al., 2025; Scheifele et al., 2012). High-intensity event music and crowd noise readily cross these boundaries and remain clearly audible in underwater recordings (Scheifele et al., 2012; Starke and Scheifele, 2008). Beyond visitor-generated airborne noise, some essential aquarium components play a massive role in habitat acoustics. Mechanical energy from life support system such as pumps generates high-intensity, low-frequency noise (typically below 100 Hz) that travels through the building’s infrastructure and propagates directly into the water (Scheifele et al., 2012; Z.- T. Wang et al., 2025). The physical characteristics of the enclosure dictate how sound behaves once it enters the tank. Because water and glass possess similar acoustic impedances, sound energy within the tank propagates easily into the glass walls (Wahlberg and Larsen, 2017). However, the interface between the outside air and the aquarium wall acts as a pressure release surface, which can cause significantly larger particle motions inside smaller aquaria than what animals would experience in the natural open waters (Akamatsu et al., 2002; Wahlberg and Larsen, 2017). Consequently, tank design heavily dictates total ambient noise levels. Our recorded noise values fall consistently within the 116.3 to 142.9 dB re 1 μPa range reported by Anderson (2013), suggesting that our study site is acoustically comparable to other captive habitats where tank construction materials and substrate composition are known to modulate ambient power levels.

## CONCLUSION

In conclusion, the investigation into teleost behavior in public aquarium reveals a complex interplay between species-specific traits, visitor total frequentation and the nature of visitor interactions. Our findings highlight that assessing visitor impact requires looking beyond public presence in front of the exhibit; instead, we must evaluate the broader environmental disturbances generated by cumulative public presence and behaviors. Furthermore, behavioral sensitivity to these visitor dynamics is highly heterogeneous across species. This divergence likely stems from differences in sensory biology—particularly species-specific auditory and visual capabilities—as well as localized coping mechanisms and micro-habitat use. Ultimately, the distinction between simple presence and public behaviors, such as tapping on glass, is critical for understanding fish welfare. The evidence suggests that while habituation is a potential long-term outcome for some, stressors can also lead to inhibitory effects that may be prevent it. Future research should prioritize integrating physiological markers, such as cortisol levels, with behavioral observations to clarify these trade-offs and test for thresholds of behavioral inhibition. These insights provide a foundational framework for optimizing design and management strategies of aquariums, thus ensuring a balance between welcoming the public and maintaining a high level of animal welfare.

## Funding/Acknowledgement

This work received support from the French government under the France 2030 investment plan, as part of the Initiative d’Excellence d’Aix-Marseille Université-A*MIDEX - Institute for Ocean Sciences (no AMX-21-IET-016). Théophile Turco was supported by the Portuguese Foundation for Science and Technology (FCT) through the strategic projects UIDB/04292/2025 (https://doi.org/10.54499/UID/04292/2025) granted to MARE, and LA/P/0069/2020 (https://doi.org/10.54499/LA/P/0069/2020) granted to the Associate Laboratory ARNET.

## Data and Code Availability Statements

The behavioral data and code used for the analyses presented in this article are available on the Zenodo repository: 10.5281/zenodo.23664809

## CRediT author statement

**Lilou Vincent**: Conceptualization, Investigation, Software, Data Curation, Validation, Visualization, Writing - Original Draft, Writing - Review & Editing; **Théophile Turco**: Methodology, Software, Data Curation, Formal analysis, Validation, Visualization, Writing - Original Draft, Writing - Review & Editing, Supervision; **Jerome Mourin**: Resources, Supervision, Writing - Review & Editing; **Joel Attia**: Conceptualization, Methodology, Validation, Resources, Writing - Review & Editing, Supervision; **Anne Sophie Tribot**: Conceptualization, Methodology, Validation, Resources, Writing - Review & Editing, Supervision, Project administration, Funding acquisition

