## Supplemental Figure for "Beyond the glass: testing visitor effects on the behavior of teleost fish in public aquaria"

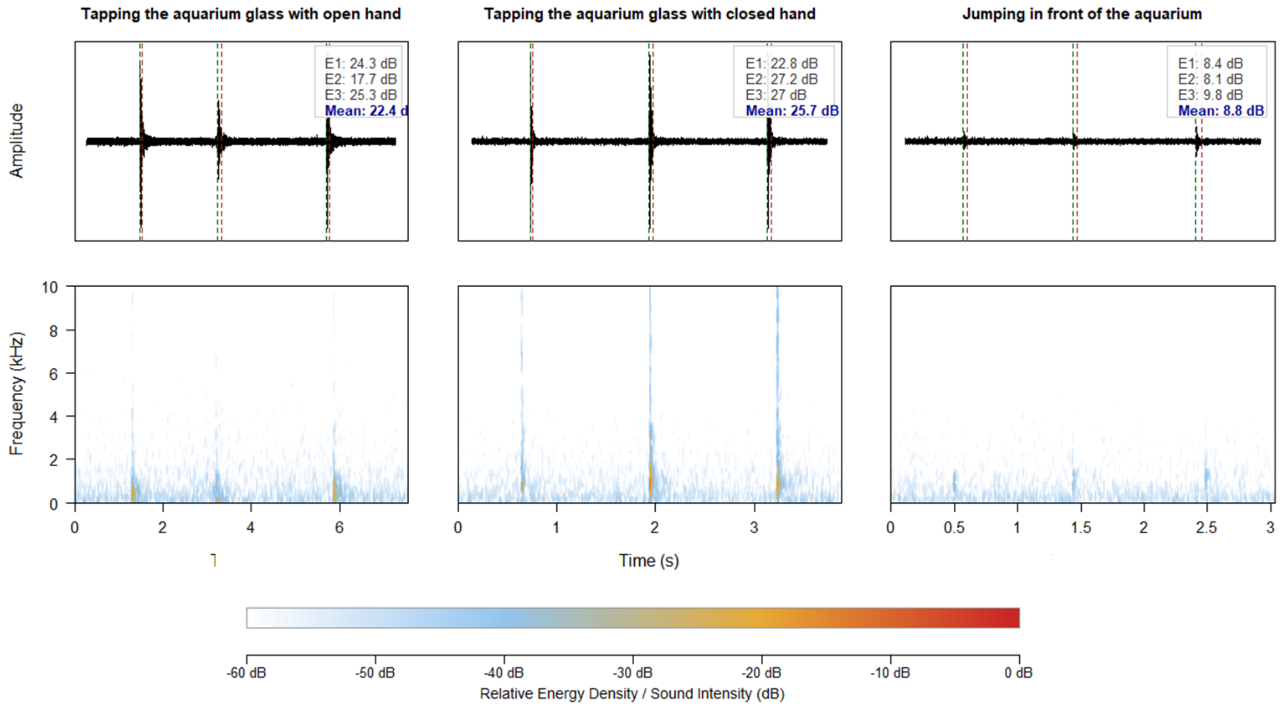


*Sup. mat. 1:* Comparative acoustic analysis of human behavior events from inside the *exhibit. The figure illustrates the acoustic profile of three conditions: tapping on the glass with an open hand (left), tapping with a closed fist (center), and jumping in front of the exhibit (right). On top, oscillograms displaying raw signal amplitude over time; vertical green and red dashed lines denote the onset and offset, respectively, of the detected acoustic events. On bottoms, spectrograms (0–10 kHz) illustrating the frequency distribution and energy density of the signals, standardized to* global *maximum amplitude across all conditions. The text boxes report the calculated Signal-to-Noise Ratio (SNR) for individual events (E1–E3) and the mean in decibels (dB). The color bar scale represents relative sound intensity in decibels (dB).*
